# Stable thalamocortical learning between medial-dorsal thalamus and cortical attractor networks captures cognitive flexibility

**DOI:** 10.1101/2021.01.15.426814

**Authors:** Siwei Qiu

## Abstract

Primates and rodents are able to continually acquire, adapt, and transfer knowledge and skill, and lead to goal-directed behavior during their lifespan. For the case when context switches slowly, animals learn via slow processes. For the case when context switches rapidly, animals learn via fast processes. We build a biologically realistic model with modules similar to a distributed computing system. Specifically, we are emphasizing the role of thalamocortical learning on a slow time scale between the prefrontal cortex (PFC) and medial dorsal thalamus (MD). Previous work [1] has already shown experimental evidence supporting classification of cell ensembles in the medial dorsal thalamus, where each class encodes a different context. However, the mechanism by which such classification is learned is not clear. In this work, we show that such learning can be self-organizing in the manner of an automaton (a distributed computing system), via a combination of Hebbian learning and homeostatic synaptic scaling. We show that in the simple case of two contexts, the network with hierarchical structure can do context-based decision making and smooth switching between different contexts. Our learning rule creates synaptic competition [2] between the thalamic cells to create winner-take-all activity. Our theory shows that the capacity of such a learning process depends on the total number of task-related hidden variables, and such a capacity is limited by system size N. We also theoretically derived the effective functional connectivity as a function of an order parameter dependent on the thalamo-cortical coupling structure.

**Significance Statement:** Animals need to adapt to dynamically changing environments and make decisions based on changing contexts. Here we propose a combination of neural circuit structure with learning mechanisms to account for such behaviors. Specifically, we built a reservoir computing network improved by a Hebbian learning rule together with a synaptic scaling learning mechanism between the prefrontal cortex and the medial-dorsal (MD) thalamus. This model shows that MD thalamus is crucial in such context-based decision making. I also make use of dynamical mean field theory to predict the effective neural circuit. Furthermore, theoretical analysis provides a prediction that the capacity of such a network increases with the network size and the total number of tasks-related latent variables.

## Introduction

In order to survive in a changing environment, animals need to acquire updated information and change plans according to the new situation and make optimal decisions on where and what to hunt [3–5]. In order to get more reward in a game like the Wisconsin card sorting task [6], participants need to flexibly change strategy and try different rules when their accuracy is dropping. This leads to a need to understand how the brain switches attention or strategy and adapts to different contexts [7, 8] by producing the appropriate behavior. This is a central effort of current neuroscience. There is only limited experimental evidence about learning mechanisms that can explain attention switching. Current research fails to explain life-long flexible learning [9], which is a critical feature drawing great attention on how neural activity in the prefrontal cortex enables animals to successfully make decisions. To date there has been limited research on how decision making depends on interactions between multiple brain regions, and how the brain makes use of the memory of prior stimuli, responses, and outcomes, to adapt to the environment. While in computer science community, the idea of distributed computing on problems like action election, leader choosing, consensus forming has already been explored, it is probably fruitful if in the neuroscience community we think more about whether the brain actually uses a distributed computing. Recently, several groups [10, 11] have tried to explore whether the brain performs parallel computing as a (k)WTA process, but communication between different brain region is more than such kind of consensus phenomenon. In this work, we will fill the gap by exploring the role of the medial-dorsal thalamus (MD) and its interaction with PFC[12, 13], which explains why PFC can do flexible cognitive switching, where functional connectivity changes according to context.

Before we talk about the role of the medial-dorsal thalamus (MD), the first question one may encounter is why the medial-dorsal thalamus? The explanation starts from the debate about whether thalamus is merely a relay hub, or it is actually doing computation based on the signal from prefrontal cortex. A recent paper [14] compared the pathway of the medial geniculate body to A1 neurons and the pathway from MD to prelimbic prefrontal cortex, and this comparison shows distinctive functions in different areas of the thalamus. The former pathway is a sensory thalamic pathway, which affects the A1 circuit mainly through excitatory neurons, so it is just a first order thalamic nucleus. For the latter pathway, MD affects the prelimbic area through inhibitory PV+ neurons, which gives modulation to the other neurons. Meanwhile, it has been reported that MD receives signals from multiple cortical areas, including the cingulate cortex (involved in hierarchical decision making), orbital-frontal cortex (involved in value-based decision making), and secondary motor cortex (involved in motor planning). This gives a structural basis for the neural computation in MD where integration of information is allowed. So, MD thalamus is a higher order thalamic nucleus, which we choose to be part of our circuit for exploring lifelong learning.

In fact, it has already been reported by multiple groups [15–18] that hypo-connectivity between MD thalamus and PFC, human or animals perform poorly in attentional or working memory tasks after training section, and tend to stick to only one rule in a previously exposed context, just as is observed in animal models of schizophrenic. Experimentalists [19, 20] have explored the role of thalamus in mice for context switching using 2 (also 4) alternative forced choice tasks. They have illuminated how the thalamus amplifies the relevant prefrontal activity while inhibits irrelevant prefrontal activity. Attention is switched by the interplay between the prefrontal cortex and the medial dorsal thalamus. Amplification is accomplished by a disinhibition pathway from thalamus to VIP interneurons and then to SOM neurons and an inhibition pathway, where the thalamus enhances the PV neuron (fast spiking neuron), and then inhibits the pyramidal neuron. However, the learning mechanism that modifies these pathways is lacking. [21] showed similar functional connectivity in humans when looking at the activity of the thalamus, orbital frontal cortex, and the vmPFC during a tactile task. They found that when the subject switches strategy, both the thalamus and orbital frontal cortex are activated, while when the subject keeps the same strategy, both the thalamus and vmPFC area are activated. Again, the detailed learning mechanism is lacking in the human tactile experiment, and it is hard to tell exactly which part of thalamus is activated due to technical difficulties. However, we do know that thalamus plays a role in changing activity in PFC. Due to the limitation of the empirical finding, theory and modeling can give more insight on possible learning mechanisms, and this will help us to design new experiments for further investigation of the learning mechanism.

Towards this end, we developed a mathematical tool to deal with the interplay between random connectivity and structured connectivity. To approach this question and understand the role of thalamo-cortical interaction in memory and learning related to context-based decision-making tasks, we focus on the role of low-rank structure of the effective coupling weight of the network. The low-rank structure in this work is mainly the thalamo-cortical connectivity weight as a building block to embed into effective weight matrix which allows the network to solve the task. Furthermore, the low-rank structure allows us to explore how exactly the neural modulation impacts the behavior of the circuit. It has already been taken as an interesting problem about how prefrontal cortex solve tasks by neural modulation [22] back in 2004. However, the theory at that time was based on the hypothesized modulation functions being multiplicative gain. Furthermore, the author specifically assumed that the factor multiplied as a context modulation term functions like a switch that enhance part of the cells while suppress the other. In this work, we will further develop the theory so that we will be able to illustrate how the plasticity and network structure of thalamo-cortical coupling led to this particular switch.

Since the thalamic units are themselves bidirectionally connected to the recurrent cortical network, there are two questions we want to address. First, how is new context information encoded into the thalamic circuitry through learning? Second, how does activity in the thalamus remap the effective connectivity of the prefrontal cortex? The first question considers plasticity in connections from the prefrontal cortex to thalamus and the second considers the reverse connections. The first allows encoding of new environments or new strategies, while the second leads to efficient information processing for adaptation to different contexts and prevents catastrophic forgetting. Here, we will explore the role of MD thalamus using low rank perturbation of the connection weight matrix in the prefrontal cortex.

Previous work mostly considered randomly connected neural networks [23–26]. More recent techniques deal with networks mixing randomness and structure focusing on motor cortex, linear networks [27] focusing on continuum movement generation and role of thalamus and basal ganglia, and nonlinear networks [28] focusing on optical control and role of thalamus modulated by basal ganglia. Work on low rank structure [29–32] exists but does not emphasize how it is related to the thalamus. To our knowledge, no one has made use of low rank structure to explore the role of thalamus in hierarchical context-based decision making, let alone its role in lifelong continual learning. This article aims to fill the gap.

Theory is helpful in giving us some conditions under which the network achieves optimal computations. We also test the theory with a specific learning. We use Hebbian learning with the additions of homeostatic synaptic scaling [33–35] and hetero-synaptic plasticity [36–40] rule. This type of learning is corresponding to the anatomy structure of thalamocortical coupling. It has been recently reported [41] that majority of the MD thalamus cells are matrix cell, meaning that they are not targeting specific region of cortex, but targeting multiple layers of cortex. This feature of matrix cell distinguishes them from the core cells, which is more likely located in sensory pathway. The core cells project to specific area of the cortex and plays the role of transmitting signal. There exists empirical evidence that both of core cell and matrix cell exist in thalamus. Since the matrix cell plays the role of modulation [42], I hypotheses that they provide the mechanism of synaptic scaling which can adjust the scale of effective coupling between MD thalamus and cortical region. There exists empirical evidence of experience-based synaptic scaling in visual system [43]. Unfortunately, no experiment has been done specifically about synaptic scaling in thalamocortical loop specifically related to MD thalamus yet. Meanwhile, I hypotheses that the same time, the core cells, which target intern-neuron, such as VIP cell, play the role of facilitating Hebbian plasticity rule. Based on these hypothesis, I make use of a network containing both random recurrent connectivity in the prefrontal cortex and a structured low rank attention module combined with thalamus, and successfully reproduce the neural dynamics observed in experiments [1]. The learning mechanism presented here meets the theoretical constraint from a mathematical analysis of dynamical mean field theory [44], eigenvalue analysis from linear algebra and random matrix theory [45]. Our framework leads to experimentally testable predictions about the population dynamics being low rank, which lead to an account of the activity of individual neurons during switches of context. Meanwhile, our theory shed light on future experiment direction, focusing on understanding the mechanism that meta-plasticity, triggered by modulation from MD thalamus, lead to synaptic scaling in PFC that allow stable and flexible cognitive function.

## Results

### A reservoir computing network [46] conducts hierarchical decision making

In a hierarchical decision-making task, the decider must first attend to either a visual or auditory attentional signal and then wait for a cue-free delay period roughly one second. Then they are presented with both visual and auditory signals and make a perception decision based on the attentional signal before the delay period. Here I emphasis that the animal only received one type of attentional signal during each trail. However, for each type of attentional signal, there are two subtypes, one represents attending to visual perception signal, and the other represent attending the auditory perception signal. The experiment [1] set up with 3 blocks, the first block and third block are presented with auditory attentional signal, while the second block is presented with visual signal. During all trials, the perception signal is the same, so animal must understand the attentional cue in order to provide the correct response, and they only get reward when they make the correct choice. The relevant training design is in Fig 2a for single trial, and Fig 2c for block. In the previous work, experimentalist found that the animal can switch between different attentional cues and make the correct decision in the perceptional decision making after the delay period. However, when they use optogenetic perturbation technology to disturb the connection between MD thalamus and prefrontal cortex, the animal can’t switch smoothly and tend to stick to the first rule they learned. The experiment then indicates that thalamus is important in the attentional selection, but not affecting the perception decision making, ruling out the possibility that the cue signal is merely a sensory input.

**Figure 1.**
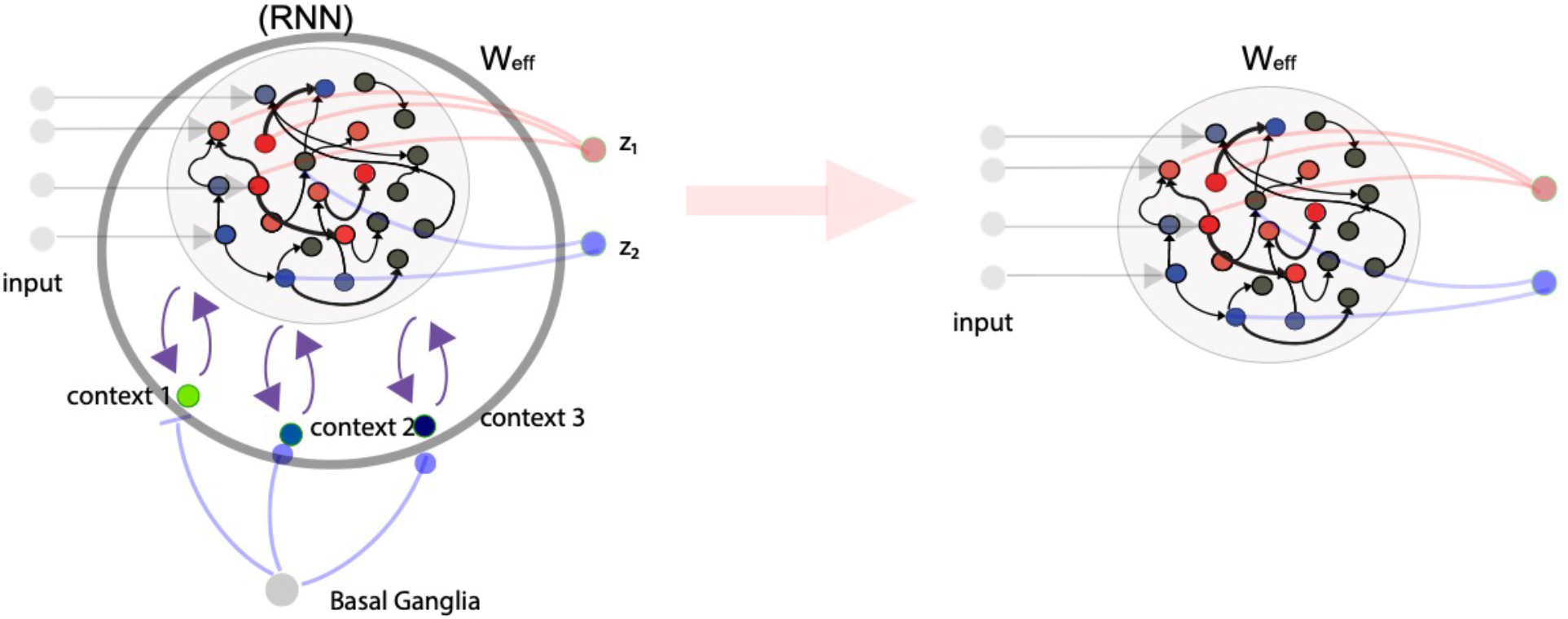
Left: Aim for showing circuit of lifelong learning, where animal should be able to switch between different context in different task and time. The two different red color neurons in PFC represent neurons which have preference to respond to sound modality, while the two different blue color represent neurons which have preference to respond the light modality. The grey color represents neurons that has no preferential to the two modalities. *z*_1_ and *z*_2_ represent two output, left and right choice. Basal ganglia cell is merely for purpose of illustration that basal ganglia will be important for more complicated tasks. Right: Also show effective weight connectivity that combine the weight within PFC and weight between PFC and thalamus. We then can compare the effective circuit and the perturbed circuit (ontogenetically perturb by reducing the connectivity between PFC and thalamus or use schizophrenia genotype animal for experiment).

**Figure 2.**
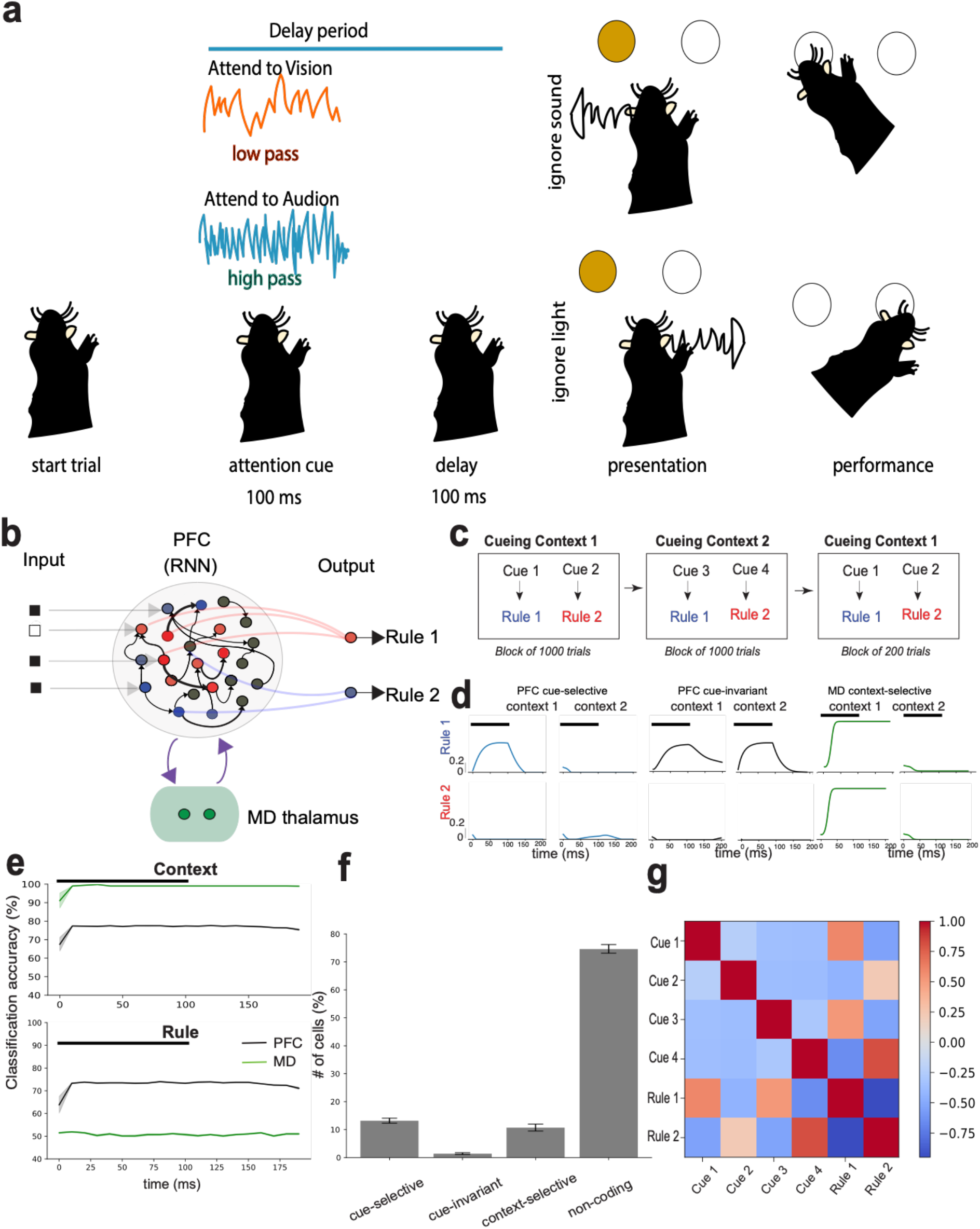
Reservoir computing model for attention task. a. The scheme plot of single trial in experiment, adapted from Fig 1 of [20]. After initial of the trial, animal is presented by attentional cue of either sound (high pass, low pass) or audition (UV and green). For simplicity, the plot only shows the case of sound attentional cue. And then based on different rule of high pass and low pass, animal need to either ignore sound perceptional signal or light perceptional signal in the presentation period. During the attentional cue, the cue lasts for 100ms, and then followed with a 100ms delay period (different from original paper where there is 400ms delay). After delay period, the perceptional signals are presented. The animal then perform choice immediately afterward. b. The architecture of the simulated hierarchy decision making network: the network includes prefrontal cortex and medial dorsal thalamus. In prefrontal cortex, neurons are recurrently coupled. However, in medial dorsal thalamus, connectivity is feed-forward. The connectivity between MD thalamus and PFC is with plasticity, with a relatively slow time scale. The connectivity between PFC neurons and output neurons are also with plasticity, with relatively fast time scale. We use the Hebbian learning rule with synaptic scaling for plasticity between MD thalamus and PFC, and we use simple mean square error (MSE) loss function for output neuron using error-correction optimization. c. The training is separated to 3 blocks. The first and third block are using context 1 (auditory input), where the attention cue is either high pass (HP) or low pass [4]. For the second block, we are using context 2 (vision input), either UV or green LED signal. The HP signal and UV signal both represent pay attention to vision, while LP and green signal both represent pay attention to audition. Animal only get reward when they do the correct respond. d. The PFC and MD thalamus firing rate activity in different input signal. We showed the cue-selective, cue-invariance and context cells. e. We do classification using support vector machine (SVM) with cross validation statistics analysis. It turns out that MD thalamus cells can distinguish the context better, while PFC cells can separate rule better. f. The bar plot for different cell types. We see that there is a lot of non-coded cells, which means most of cells in PFC contain mixed information. g. The correlation relation between different cell groups. We see that the PFC cells selectively responding to 4 cues are mainly enhancing activity of each other, while suppressing the other cue activity. The PFC cells corresponding to rule 1 is strongly correlated to the PFC cells activity of cue 1 and cue 3, while those corresponding to rule 2 is strongly correlated to PFC activity of cue 2 and cue 4. This is consistent with other evidence in our simulation.

In order to model the task explained above, we consider a randomly connected attractor network with N neurons, labeled *i*, each with synaptic input, *I*_*i*_, which varies according to

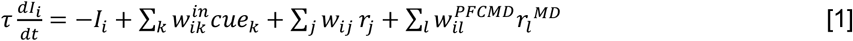

where 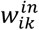 is the input weight, *w*_*ij*_ is the weight between neuron *i* and *j* in PFC, 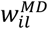 is the coupling weight between neuron *i* in PFC and neuron *l* in MD thalamus.

The circuit is shown in Fig 2b. We conjecture that there exists a genetically hard-wired motif in prefrontal cortex, where a particular type of input is acted on. This is reasonable because inputs corresponding to different contexts are from different cortex regions, similar to the model in recent work [47]. The input passes through the activation function:

*r*_*j*_ = tanh *I*_*j*_ if *I*_*j*_ > 0, otherwise *r*_*j*_ = 0

The outputs from the network obey

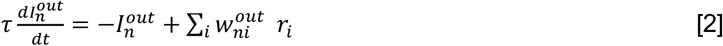

where
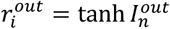 if 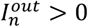, otherwise 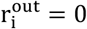

The weights update according to the rules

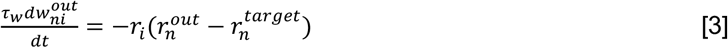

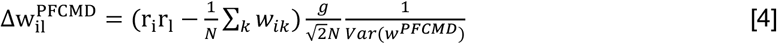

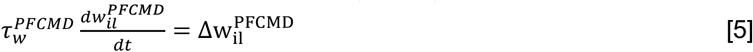

The 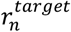 is the correct response for the *n*-th output neuron. The 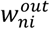 is the weight from i-th PFC neuron projecting to the n-th output neuron. The update of w_il_ contain the Hebbian term r_i_r_l_ and the synaptic scaling term 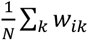, where we are trying to keep 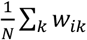 as constant. We normalize the weight using factor 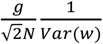 so that the total variance of the weight remains bounded.

The network was trained based on the protocol in Fig 2c. With same scheme as other reservoir computing networks, we keep the input weight and coupling in PFC as fixed weight, but the thalamo-cortical weight and the output weight are being trained. The former weight changes with a slow portion in a trial, while the latter weight is changes in a trial. Based on their task-dependent activity, I identified cue-selective and rule-selective units in the PFC, and the example cells are presented in Fig 2d. Applying the support vector machine (SVM) using all trials for each time step within a trial, PFC cells are found to be able to classify the different rules, while media-dorsal (MD) thalamic cells do not. I also find that MD thalamic cells store more information about the context. I then get the statistics of 4 different cells in PFC, cue-selective cells, cue-invariant cells, context selective cell and non-coding cells. I find that the majority of cells are mixed encoding cells. I also show the correlation in Fig 2g.

### Activation of thalamic cells enhance relevant activity in PFC while suppressing irrelevant activity

To further understand the switching, I show the transient dynamics during switching for both thalamic cells and PFC cells. In experiment [1], using support vector machine, it is observed that MD thalamus cells can be classify context information with statistical significance. Based on the experimental design, in the first and third block, the activity of cells which are cue-selective specifically to cue 1 and cue 2 are relevant response, while the activity of cells which are cue-selective specifically to cue 3 and cue 4 are irrelevant response. Previous work [1] made use of the neural data and GLM method for analysis, and the found that the MD thalamus activity is positively correlated to neural response of cells selective to the cue 1 and cue 2, but negative correlated to the neural response of cells selective to cue 3 and cue 4 in the first block. In our simulation, we reproduced the findings in experiment. Fig 3b shows that the first MD thalamic cell was randomly selected as encoding context 1 by the network. Context 2 is encoded by the second context cell. This is consistent with the experimental observation that MD thalamic cells can encode contexts and MD thalamus cell can distinguish different contexts if the contexts are switching slowing. We also get the weight dynamics over trials in Fig 3g, which matches experimental observation that the thalamic cells enhance relevant response and suppress irrelevant response.

**Figure 3.**
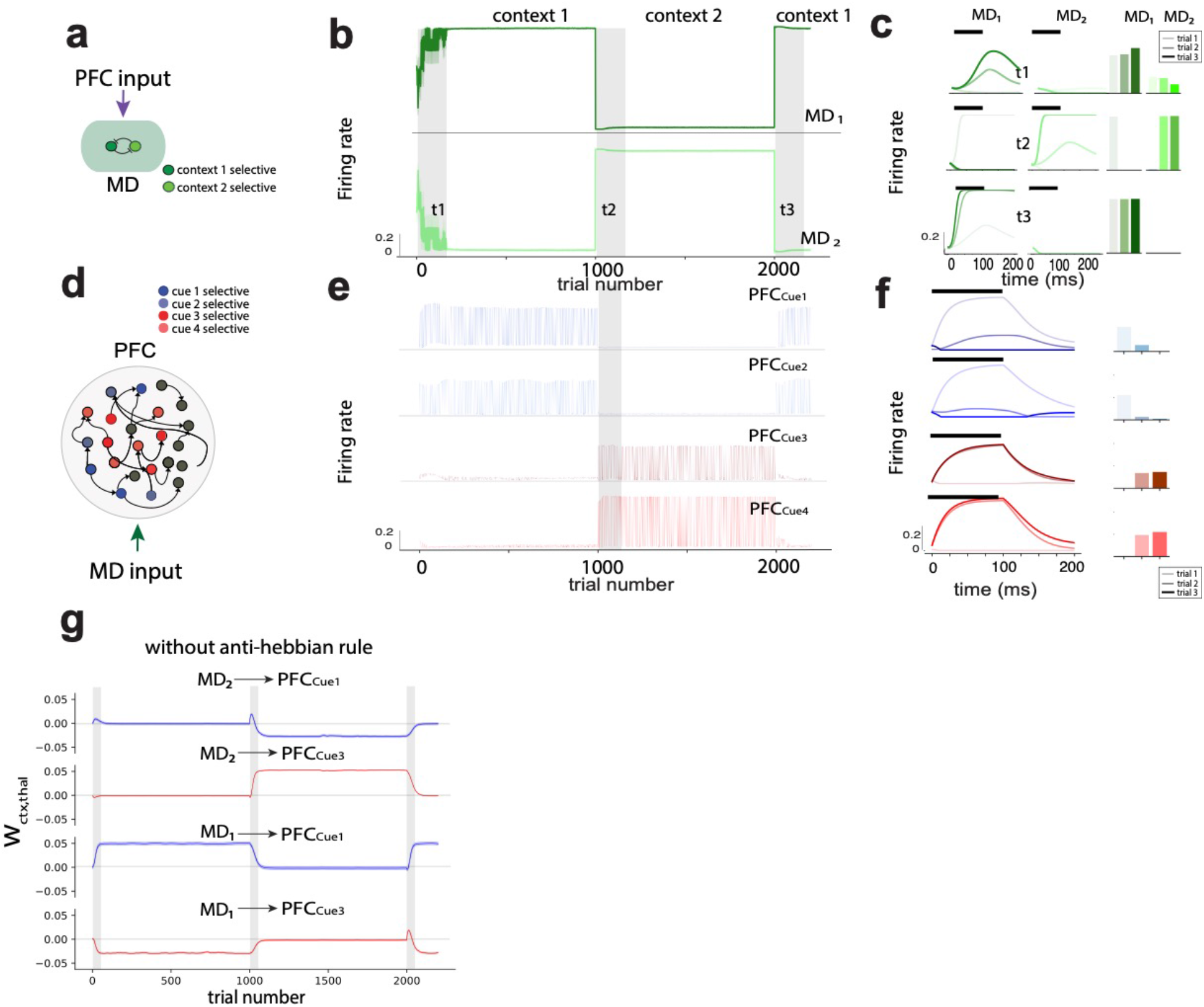
Learning mechanism with medial dorsal (MD) thalamus cell a. Schematic plot for medial-dorsal (MD) thalamus cell circuit. It mainly receives excitatory input from prefrontal cortex (PFC), but it also receives inhibitory input from other thalamus cells. The circuit then become winner-take-all circuit effectively, similar to leader-selection problem in computer system. b. The firing rate activity of the two MD thalamus in simulation. The upper cell only active during context 1, while the lower cell only active during context 2. c. The cell activity during trials period t1, t2 and t3. During t1, the first MD thalamus cell gradually increasing activity, while the second MD cell are getting suppressed. During t2, the first cell gradually decrease activity, while the second cell gradually decrease activity. In t3, the first cell increase activity very quickly, while the second cell get suppressed to silent almost immediately. d. The schematic plot for prefrontal cortex cue selective cell activity. e. The firing rate activity of the prefrontal cortex cells selectively respond to the 4 different cues. We see that those selectively respond to cue 1 and cue 2 only get activated during context 1, and those selectively respond to cue 3 and cue 4only get activated during context 2. f. The firing rate activity of the 4 type of prefrontal cortex cells during trials t2. g. The coupling weight evolution plot during all trials. During context 1, MD 1 cell is enhancing activity of PFC cells selectively respond to cue 1 but suppressing activity of PFC cells selectively respond to cue 3. However, during context 1, MD 2 cell keep the coupling to these PFC cells to be zero. In context 2, MD 2 cell is enhancing PFC activity selectively respond to cue 3 but suppressing PFC activity selectively respond to cue 1. And in context 2, MD 2 cells is keeping zero connectivity to those PFC cells. This result is key result where we get evidence that thalamus cells control the PFC activity by enhancing the activity related to the current context, while suppressing the activity irrelevant to current context. This allows flexible (fast forget old information and fast learn new information) cognitive functions.

### Slow change in thalamic connectivity to PFC does not affect PFC activity in response to fast changes of context

To test the role of fast changing of output neuron and slow changing of the thalamo-cortical reciprocal connectivity, we simulate the case where context is changing frequently instead of simulating the block setting shown in Fig 2. We find that with frequently changing context, MD thalamic units assign all 4 different inputs into one context, and the animal’s decision is mainly determined by the training of the output neuron. The resulting firing rate dynamics of MD thalamus is shown in Fig 4a. In Fig 4b, we show the mean square error of the output neuron, and it turns out that it keeps optimizing and it does not show a peak when context changed. The result matches the experimental finding in ([19]).

**Figure 4.**
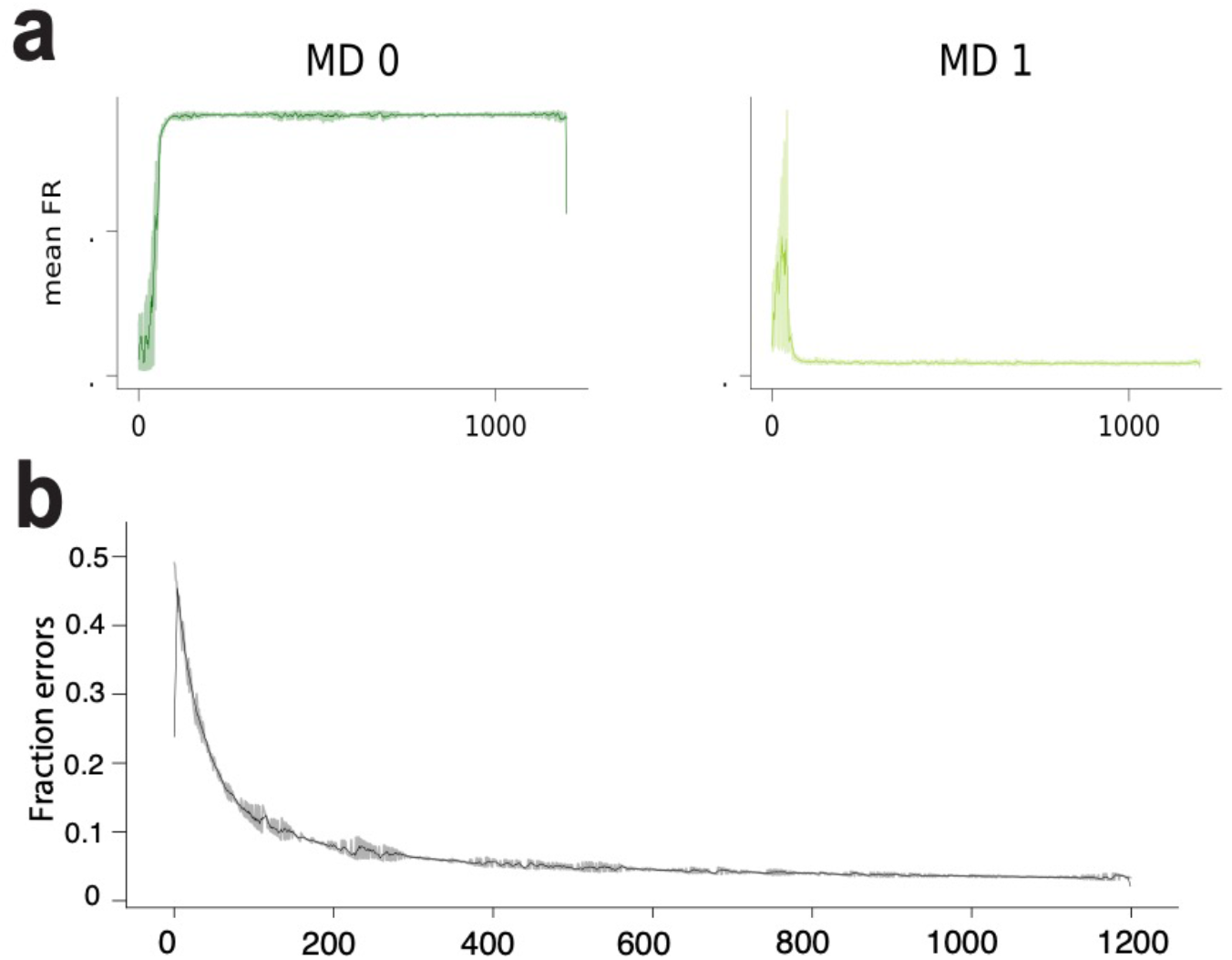
Fast switching of context, plasticity between output neuron and prefrontal cortex neuron plays main role. a. Activity of the 2 medial-dorsal (MD) thalamus cells. We see that although the context is frequently switching, the two cells only encode on context. This means that in this setting, the slow learning between MD and PFC don’t play a role in correct respond of PFC cells. b. The mean square error of the output neuron. The network is just keep optimizing the output neuron, which his fast learning. This means when context is switching fast, it is key to associate the cue selective PFC cells to the output neuron. This also shows that the neural circuit is doing different learning mechanism for different context changing frequency. The slow learning between PFC and MD thalamus only play a role when context is changing relatively slow.

### MD thalamus disconnection results in catastrophic forgetting

To test the model further, we perturbed the network by introducing a lesion of the MD thalamic cells. The result is shown in Figure 5, where 5a is the schematic plot for the network under MD disconnection, while 5b is the network’s performance. In 5b, the upper one is for the full model, while the bottom one is for MD disconnection. For intact MD, there is less catastrophic forgetting, and this shows that the network can switch flexibly between tasks when the same context is shown again, while without MD, there is a bigger mean square error for the output, meaning a disruption of flexible switching.

**Figure 5.**
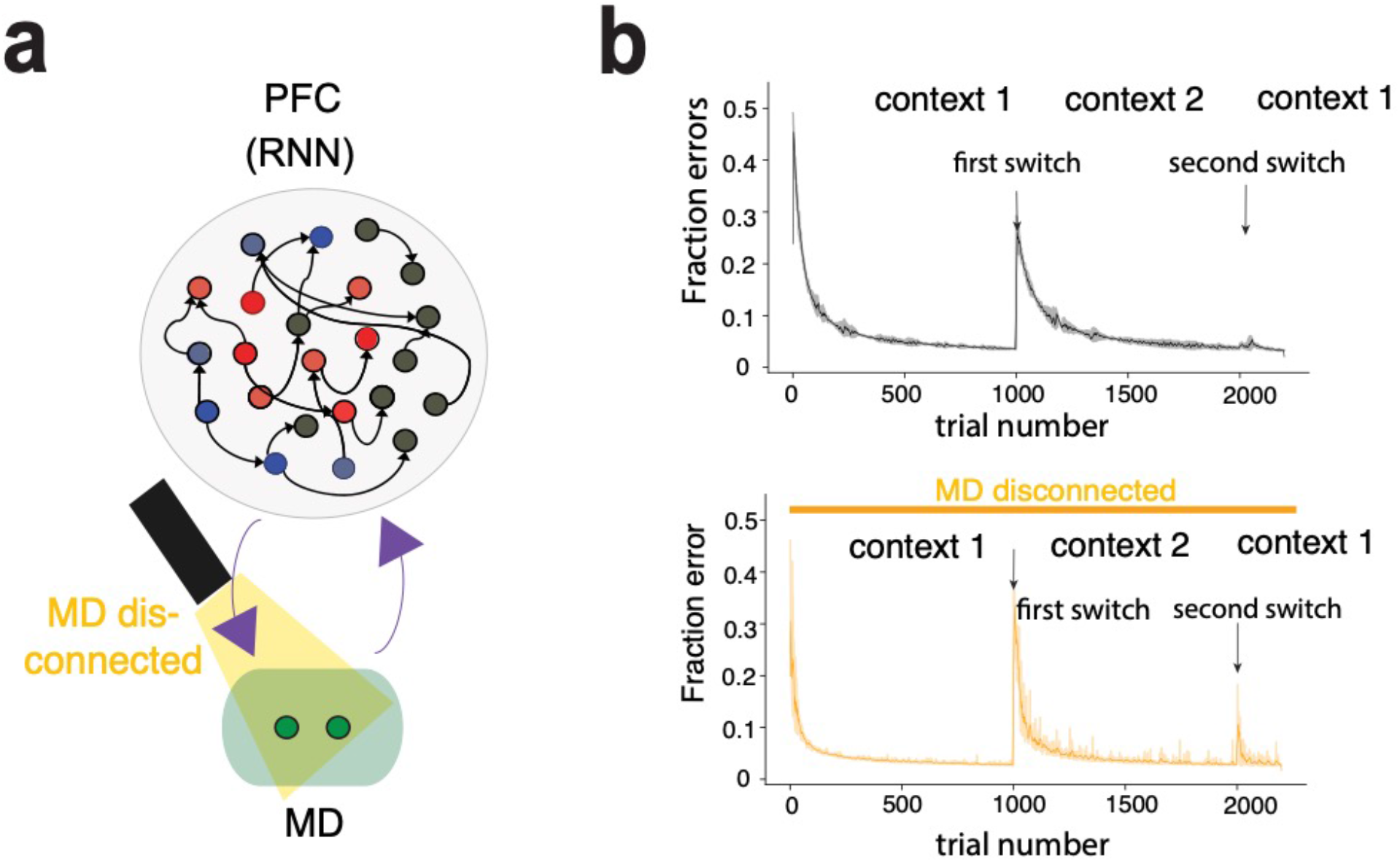
catastrophic forgetting become obvious during MD disconnected trials. a. The circuit schematic plot for MD disconnection, which is mimicking the experimental setting using optogenetic perturbation to activity of MD thalamus cells. b. Comparison between the mean square error of full model and model with MD disconnected. The black one is for the full model, and we see that during second switch of context (from context 2 to context 1), the circuit can keep doing correct choice without much error. But when MD thalamus is disconnected, showing in the yellow curve in the bottom, the network needs to relearn the context 1 and a relatively large peak is shown during the second switch. This means adding thalamus in the circuit can avoid catastrophic forgetting phenomenon, a phenomenon that catching attention in machine learning community for a long time.

## Discussion

In this work, we combined the tools of computational simulation and theoretical analysis to explore context-based decision making. Distinct from traditional decision-making models where the context input is just a separate set of inputs, we emphasize the role of medial-dorsal (MD) thalamus on neural modulation to the prefrontal cortex, which is crucial factor that introduce cognitive flexibility. This mechanism allows attentional control of the neural response to the context cue signal and facilitate efficient perceptional decision-making after delay period. We also notice there is empirical evidence showing neural modulation of functional cortical connectivity due to thalamic activity [48]. We emphasize that learning in the neural circuit can arise purely from changes in the thalamo-cortical coupling weight. Similar to traditional work on context-based decision making by Fusi et. al. [49], we found a different connectivity pattern and therefore neural response pattern when context information is switching fast, compared to the case where context information is relatively stable and changes slowly. With the simulation data, we use support vector machine to classify the firing rate data of all trials, and we observed better classification on rules in PFC cells, while better classification of context in thalamus cells, just as observed in the analysis of experimental data. We also observed different type of cells, including cue-selective cell, cue-invariance cell, context selective cell and mix-coding cell. Based on different cell type, we analyze the correlation between different cell type and see strong correlation between context selective cell and the corresponding cue cells. Furthermore, during learning, we see suppression of irrelevant cue-selective cell from thalamus and enhancement of relevant cue-selective cell, which verified the role of modulation of MD thalamus. In order to more deeply understand the effect of thalamo-cortical learning, we theoretically verified that with modulation of PFC activity arising from a structural perturbation of the thalamo-cortical architecture, we are able to gain cognitive flexibility because different thalamic activity states result in different effective circuits in the PFC. We also test the model by disconnecting the MD thalamus and check the performance of learning. We found that when MD thalamus is disconnected, the learning suffers from catastrophic forgetting. This is consistent with experimental discovery. In summary, by statistical analysis and theoretical analysis, we verified that our neural network with Hebbian learning rule modulated by synaptic scaling is biological plausible. We also further use analysis of Jacobian of the equation of motion and figured that the network can embed multiple dynamics patterns as long as the total amount of latent variable is less than system size. Combining this observation with the fact that our learning rule can result in distinct attentional state in MD thalamus, we figured that thalamo-cortical plasticity rule can facilitate transition of attractor state between different attentional state, which remains to be tested by more powerful model with larger thalamus population in the future.

### Extension and future direction

From a theoretical perspective, recent ideas from machine learning and meta learning [50, 51] provide us an opportunity to extend the model. We can apply a similar analysis with different algorithms in training and using a different model of plasticity, including the back propagation through time (BPTT) algorithm for training and compare the results with those of the current algorithm. There is an argument that reservoir computing cannot store information of time when the network reach a fixed point([52]), so it will be a good update if we can figure out how BPTT algorithm (with RNN) will help solving context dependent task with more timing information included. This is potentially more helpful for accumulation tasks [53, 54]. One of the ways to deal with this problem is introducing decision-making model that include ramping activity, which will show some time dependency in the process.

We can apply our method to animal model of schizophrenia and see whether we can understand the reason for loss of performance when an animal model of schizophrenia is engaged in various tasks [55]. Generally, it is claimed that the way schizophrenia animal got malfunction is due to the reduction in strength of the thalamo-cortical connectivity[56, 57]. Our simulation result is supporting this statement. We then probably can expend the similar modeling method to analyze sensory processing that is related to Autism, especially in visual system. Since the plasticity in thalamocortical (LGN and pulvinar nuclei will be two possible related thalamus areas) connectivity provide the modulation to PFC and adjust the E-I balance, it will be interesting to see whether our model can explain behavior deficit in Autism model of animal.

We can apply this method to the auditory system, where the relative timing of inputs is very important. We may need to add synaptic depression because the medial geniculate body (MGB) in auditory pathway probably responds on a faster time scale, which results in difficulty in affecting activity in PFC. It will be interesting to explore whether integrating input from other brain region will modify the neural response of auditory input, which is probably related to relay of different nuclei of thalamus. However, recent research shows that synaptic depression in PFC alone is more important than the feedforward inhabitation from MGB, making it questionable about the role of thalamus. Further investigation on this will be beneficial.

It will be interesting to link the relation between thalamo-cortical architecture and the attention module as incorporated in the artificial intelligence field. However, one can also integrate the effect of thalamo-cortical plasticity into the plasticity rule within PFC, resulting in seemingly homogeneous network. This leads to debate between experimentalist and theorist, even between different community of theorists about how to build a model that is biologically plausible. Theoretical effort is needed to close this debate and show that the two type of network is equivalent.

## Materials and Methods

### Network model and learning rule

We consider a randomly connected attractor network with N neurons, labeled *i*, each with synaptic input, *I*_*i*_, which varies according to

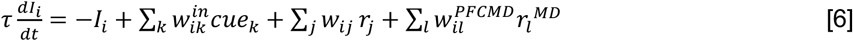

where 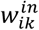 is the input weight, *w*_*ij*_ is the weight between neuron *i* and *j* in PFC, 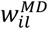 is the coupling weight between neuron *i* in PFC and neuron *l* in MD thalamus.

The total synaptic input passes through the activation function:

*r*_*j*_ = tanh *I*_*j*_ if *I*_*j*_ > 0, otherwise *r*_*j*_ = 0

The outputs from the network obey

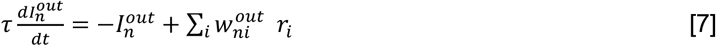

where
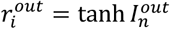 if 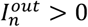, otherwise 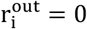

The weights update according to the rules

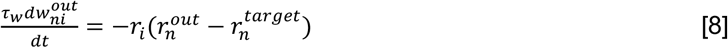

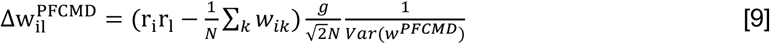

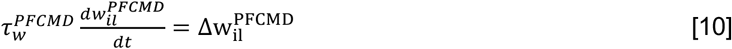

The 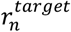 is the correct response for the *n*-th output neuron. The 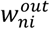 is the weight from i-th PFC neuron projecting to the n-th output neuron. The update of w_il_ contain the Hebbian term r_i_r_l_ and the synaptic scaling term 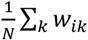, where we are trying to keep 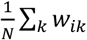 as constant. We normalize the weight using factor 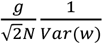 so that the total variance of the weight remains bounded.

### Simulation and analysis

All simulations and analysis are using python packages (Python 3.7, Pytorch for GPU). The simulation of rate model is using simple Euler method with time step *h* = 0.01. The low rank reduced model is in SI Apendix, where the simulation of possible dynamics is solving simple ordinary differential equations (ODE) using Euler method. Using low rank analysis, we get the effective circuits for different context and produces different neural responses for different context. The relevant analysis is shown in Fig 6, 7 and 8. The analytical analysis reveal that neural modulation is the main mechanism that enable attentional selection, where thalamus influence the functional connectivity within PFC circuit through modulation of the gain function by changing coverability. This is different from traditional view of attentional signal is just a simple input transmitted by thalamus.

**Figure 6.**
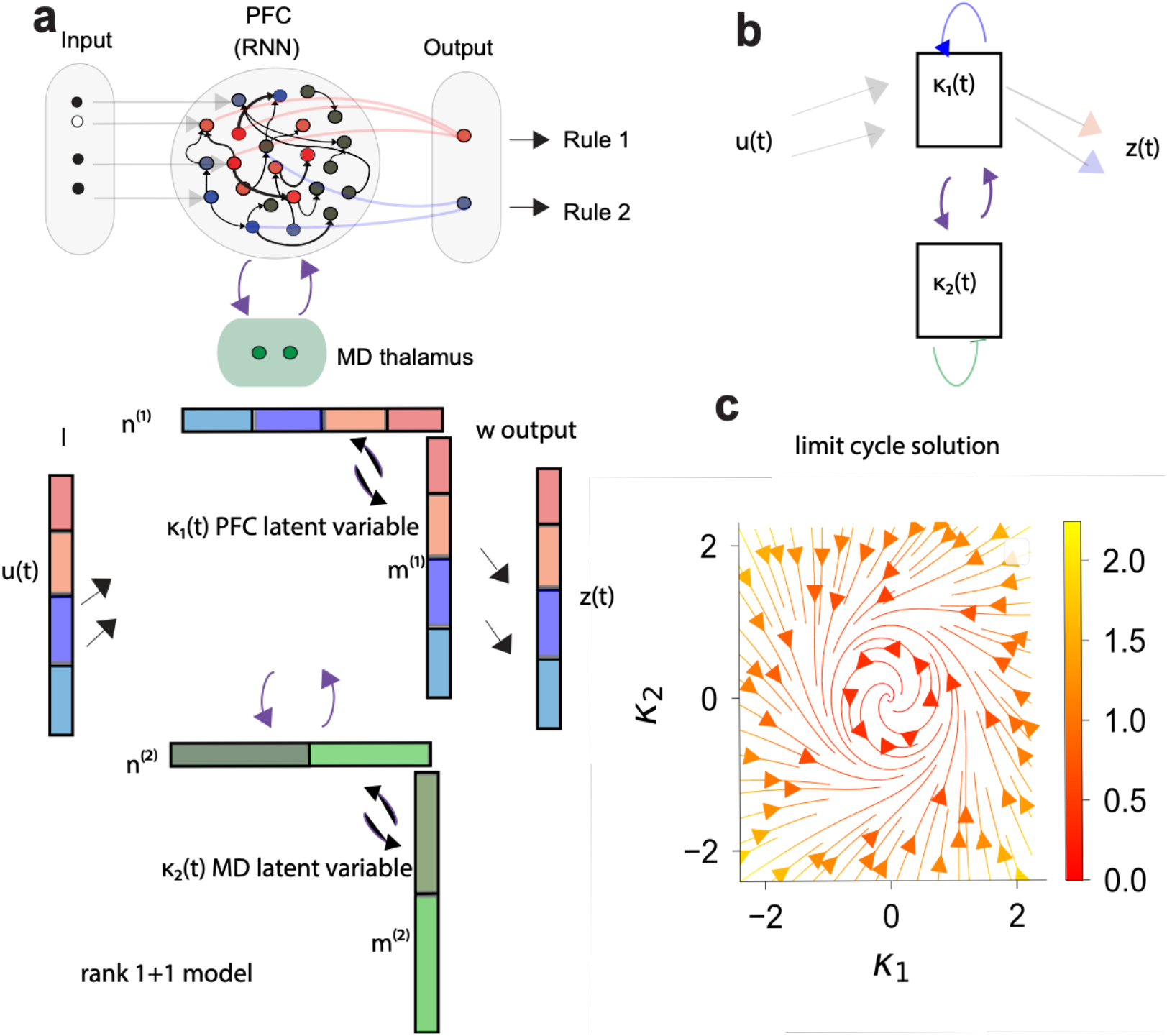
2D rank, multiple population model a. The original full model schematic plot on the top, and multiple population low rank reduction model schematic plot on the bottom. b. The effective model schematic plot, u(t) is the input, *κ*_1_ and *κ*_2_ are the two mean field variable representing PFC and MD. *z*(*t*) is the output. c. With different parameter setting of the covariance between different component (m and n in our case) in the reduced model, network can produce arbitrary dynamics. This is the example of the right parameter for creating a limit cycle activity.

**Figure 7.**
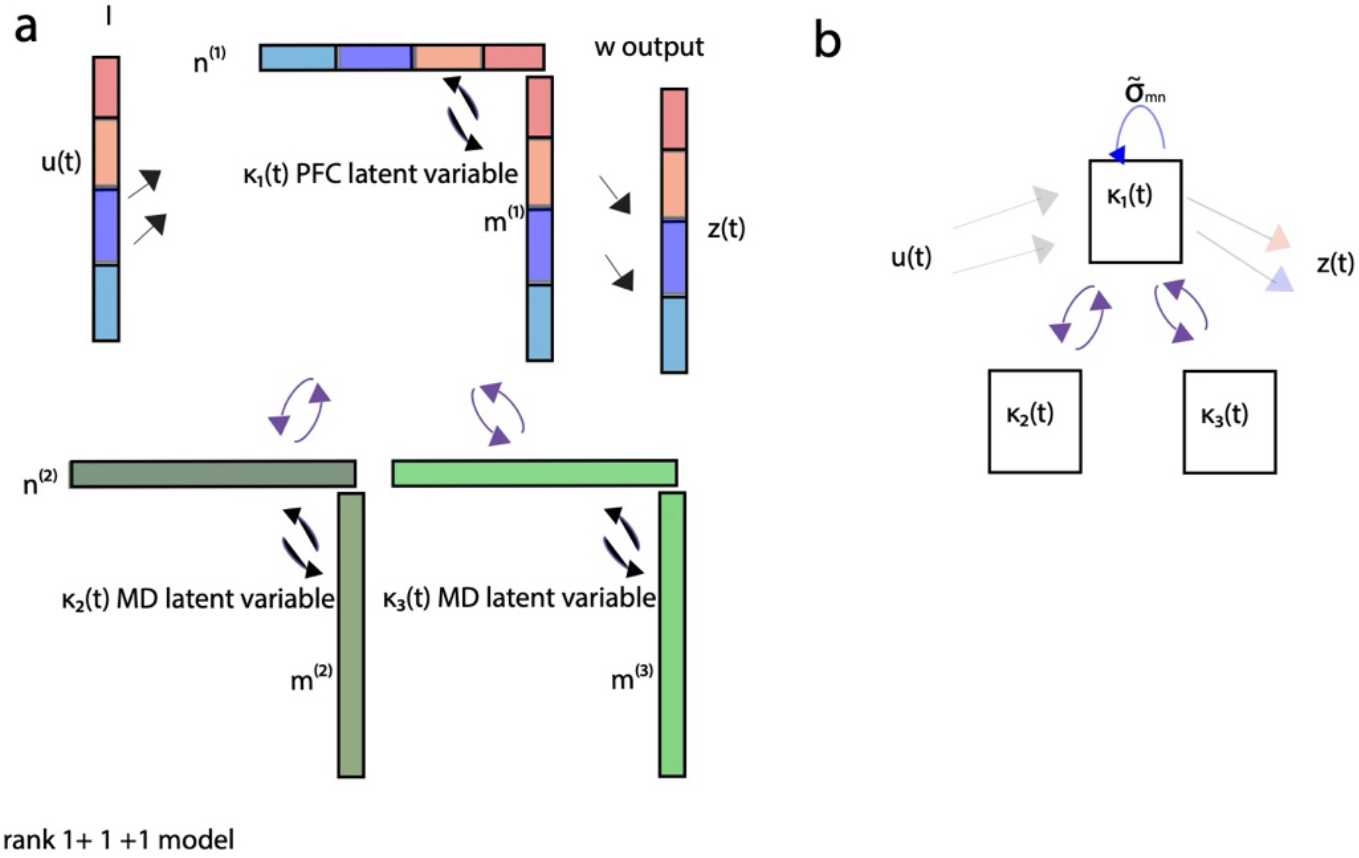
The schematic plot for model with 2 MD cells a. The reduced model scheme plot b. The effective circuit we get from this model.

**Figure 8.**
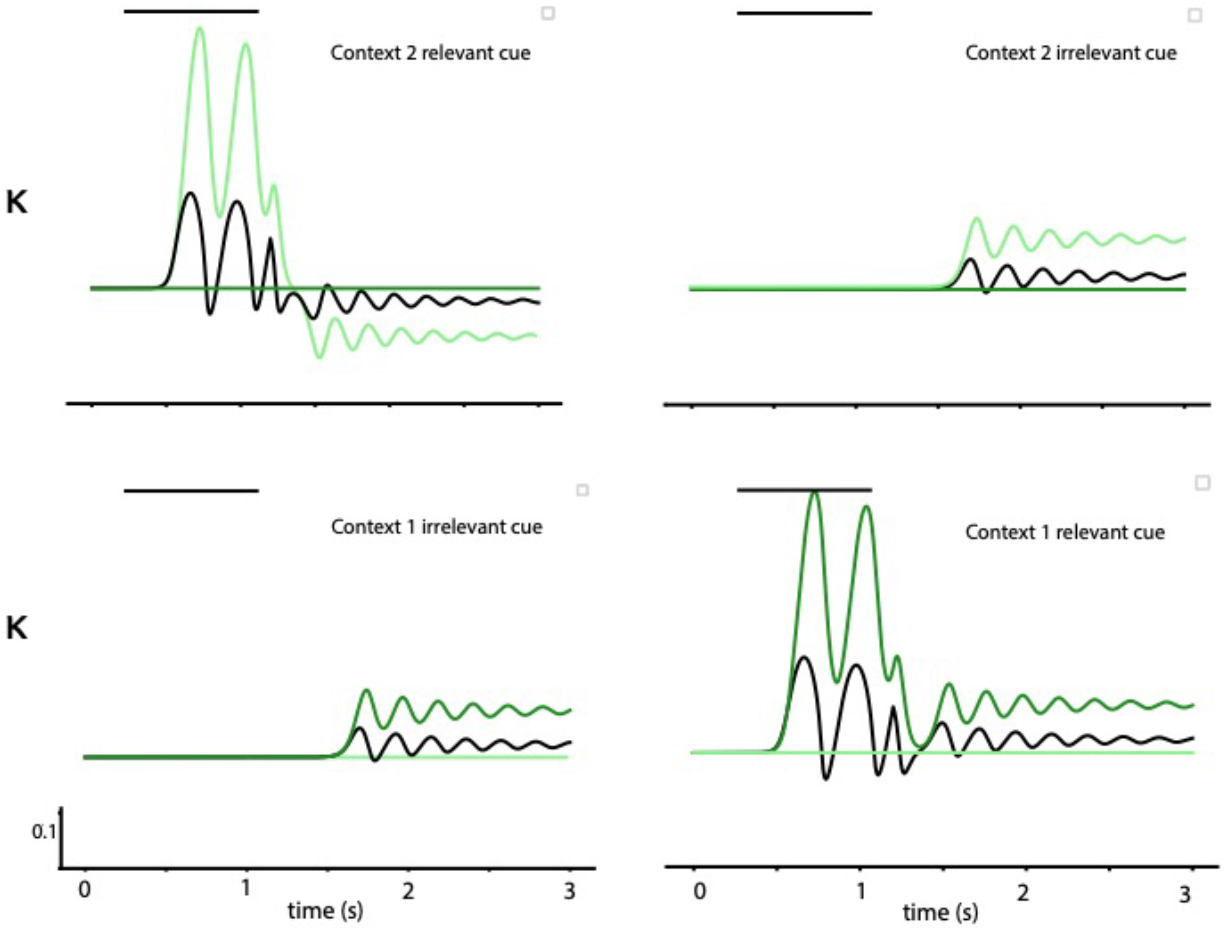
Context dependent computing by modulating the gain The upper two plots are corresponding to context 2, while the lower 2 plots are corresponding to context. We compare the response to relevant cue and irrelevant cue. And we show that the network only responds to the relevant cue.

## Data Availability

In this work, only simulation data is available, it will be available in Github at https://github.com/qius5/Mouse_paper once the paper is published.

## Acknowledgments

We are grateful to thank Michael Halassa and other people in MIT group for the discussion about his experimental findings on the early stage of this project. We thank Paul Miller in Brandeis University and Carson Chow in NIH for deep discussion on the theory related to synaptic scaling and proof-reading of the manuscript. This work is funded by Eve Marder’s training grant NIH T32NS007292.

---

**Table 1.** Overview of key quantities of reservoir computing network

| Table 1. Overview of key quantities of reservoir computing network |  |
| --- | --- |
| $N$ | Number of units |
| $I_i$ | Total synaptic input of neuron $i$ |
| $w_{ij}^{in}$ | Input weight, fixed weight |
| $w_{ij}$ | Coupling weight within PFC, fixed weight. |
| $w_{ij}^{PFCMD}$ | Coupling weight between MD neuron $j$ and PFC neuron $i$, with plasticity |
| $r_i$ | Firing rate of PFC neuron $i$ |
| $r_i^{MD}$ | Firing rate of MD neuron $i$ |
| $r^{out}$ | Firing rate of output neuron |
| $r^{target}$ | Correct respond 1, or 0(fire or silent) |
| $\tau_w$ | Time constant of output weight |
| $\tau_w^{PFCMD}$ | Time constant of thalamo-cortical weight |

