## Supplemental text for "Stable thalamocortical learning between medial-dorsal thalamus and cortical attractor networks captures cognitive flexibility"

### Appendix

We describe the low rank analysis for dynamical mean field theory in more detail. And then we give some derivation to show the capacity of the network to encode multiple contexts depends on network size $N$ and the total number of the task variables.

**Low rank dynamical mean field theory for thalamo-cortical structure**

If the structure of network meets the following conditions:

(1) It includes a recurrent network, which is randomly connected with weight as the random variable obeying Gaussian distribution $\mathcal{N}(0, \frac{1}{\sqrt{N}})$.
(2) It includes an attention module, which is composed of two thalamic cells
(3) The recurrent network and attention module are bi-directionally connected, and coupling is a matrix $W^{MD}$.
(4) Transfer function is tanh

we then can construct an effective network whose coupling weight is the sum of a gaussian random component ($\chi_{ij} \in \mathcal{N}\left(0, \frac{1}{\sqrt{N}}\right)$) and structured components ($P_{ij}^1$ and $P_{ij}^2$):

$$W_{ij} = g\chi_{ij} + P_{ij}^1 + P_{ij}^2 + O(\frac{1}{N})$$

where $P_{ij}^l$ has the form $\frac{1}{N} W_{il}^{MD} W_{lj}^{MD}$ , which is an outer product $uv^T$for vectors $u$ and $v$.

and we have normalized by N so that each coupling weight is of order $\frac{1}{\sqrt{N}}$ and $\chi_{ij}$ is a gaussian variable with standard deviation $\frac{1}{\sqrt{N}}$.

Derivation:

Thalamus has the following rate dynamics:

$$r_l^{MD} = \tanh\left(I_{PFC\to MD}\right)$$

$$I_{PFC\to MD} = \frac{1}{\sqrt{N}} \sum_j W_{lj}^{MD} \phi(I_j)$$

For transfer function $\phi(x) = \tanh x$, one has $\phi(x) \approx x - \frac{x^3}{3}$, one has:

$$r_l^{MD} \approx \frac{1}{\sqrt{N}} \sum_j W_{lj}^{MD} \phi(I_j) - \frac{1}{3N^{\frac{3}{2}}} \left( \sum_j W_{lj}^{MD} \phi(I_j) \right)^3$$

We then get:

$$\frac{1}{\sqrt{N}} \sum_j W_{ij}\, r_j + \frac{1}{\sqrt{N}} \sum_l W_{il}^{MD} r_l{}^{MD} = \frac{1}{\sqrt{N}} \sum_j W_{ij} \phi(I_j) + \frac{1}{N} \sum_l W_{il}^{MD} W_{lj}^{MD}\, \phi(I_j) + O(\frac{1}{N^2})$$

We can write down the recurrent sum in the form of $\frac{1}{\sqrt{N}} \sum_j W_{ij}^{eff} \phi(I_j)$, where we have:

$$W_{ij}^{eff} = W_{ij} + \frac{1}{\sqrt{N}} \sum_l W_{il}^{MD} W_{lj}^{MD}$$

We show that we have effective theory where effective coupling weight is well defined under large $N$ limit. The leading order contribution is of low rank $l \ll N$.

For convenient, we denote $m_i^{(l)} = W_{il}^{MD}$, while $n_i^{(l)} = W_{li}^{MD}$

Following paper [1] In large $N$ limit, for equilibrium state, mean of total synaptic input has the following form:

$$\mu_i = \sum_l \kappa_l m_i^{(l)} + I_{i,cue}$$

$$\kappa_l = \frac{1}{N} \sum_{j=1}^{N} n_j^{(l)} [\phi_j]$$

Variance of total synaptic input is the following form:

$$\Delta_i = g^2 \langle [\phi_i^2] \rangle$$

Following paper [1], we can have the following closed form of equation set for a rank 1 model for stationary solution:

$$\mu := \langle [x_i] \rangle = \langle m_i \rangle \kappa$$

$$\Delta_0 = \langle [x_i^2] \rangle - \langle [x_i] \rangle^2 = g^2 \langle [\phi_i^2] \rangle + (\langle m_i^2 \rangle - \langle m_i \rangle^2)\kappa$$

$$\kappa = \int dm \int dn\, p(m,n)\, n \int \mathcal{D}z \phi(m_i \kappa + \sqrt{\Delta_0^1} z)$$

$$\langle [\phi_i^2] \rangle = \int dm\, p(m) \int \mathcal{D}z \phi^2(m_i \kappa + \sqrt{\Delta_0^1} z)$$

$$p(m,n) = \frac{1}{N} \sum_{j=1}^{N} \delta(m - m_j)\delta(n - n_j)$$

Furthermore, if one parameterizes the problem using $M_m$ and $\Sigma_m$ as mean and variance of vector $m$, while using $\rho$ as correlation, defined by $\rho = \frac{\langle m_i n_i \rangle - M_m M_n}{\Sigma_m \Sigma_n}$, we have:

$$m = M_m + \Sigma_m \sqrt{1-\rho} x_1 + \Sigma_m \sqrt{\rho} y$$

$$n = M_n + \Sigma_n \sqrt{1-\rho} x_2 + \Sigma_n \sqrt{\rho} y$$

$$\mu = M_m \kappa$$

$$\Delta_0 = g^2 \langle [\phi_i^2] \rangle + \Sigma_m^2 \kappa^2$$

$$\kappa = M_n \langle [\phi_i] \rangle + \kappa \rho \Sigma_m \Sigma_n \langle [\phi_i'] \rangle$$

We are also interested in the stability matrix, which is summarized as follows:

$$\mathcal{M} = \begin{pmatrix} 0 & 0 & M_m \\ 2g^2 \langle [\phi_i \phi_i'] \rangle & g^2 \{ \langle [\phi_i'^2] \rangle + \langle [\phi_i \phi_i''] \rangle \} & 2\Sigma_m^2 \kappa^0 \\ 2bg^2 \langle [\phi_i \phi_i'] \rangle & bg^2 \{ \langle [\phi_i'^2] \rangle + \langle [\phi_i \phi_i''] \rangle \} & 2\mathrm{b}\Sigma_m^2 \kappa^0 + a \end{pmatrix}$$

Here, $a$ and $b$ are defined as follows:

$$a = \langle M_m M_n + \rho \Sigma_m \Sigma_n \rangle \langle [\phi_i'] \rangle + \rho \kappa^0 M_m \Sigma_m \Sigma_n \langle [\phi_i'''] \rangle$$

$$b = \frac{1}{2} \{ M_n \langle [\phi_i''] \rangle + \rho \kappa^0 \Sigma_m \Sigma_n \langle [\phi_i'''] \rangle \}$$

In order to get the low dimensional representation of the system, we use singular value decomposition ([2]). In any singular value decomposition $M = U \Sigma V^*$, one can have $M = \sum_i \sigma_i U_i \otimes V_i$, where $\otimes$ is outer product.

By doing so, we are able to interpret the effective coupling strength as the functional connectivity between two different brain regions. Furthermore, the prefrontal cortex is receiving the neural modulation coming from the structured connectivity between the PFC and the thalamus. This is due to the geometric relation between the coupling weight eigenvectors and the input vectors.

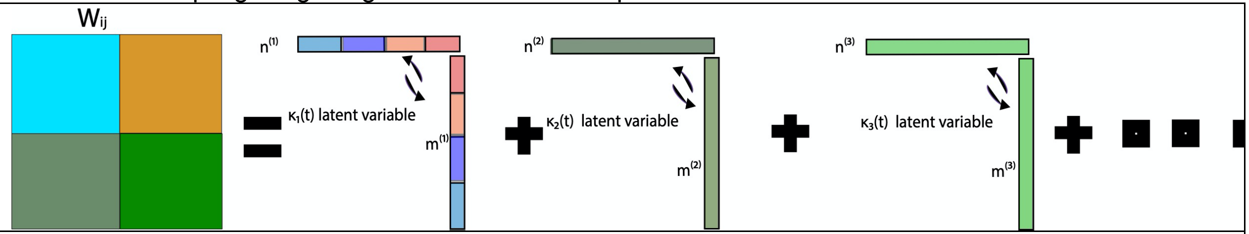


Figure S1: A schematic of the basic idea of singular value decomposition.
This shows that by doing such a decomposition, each task-related variable can be one term. We then can embed the effective connectivity of prefrontal cortex with multiple task-related variables, allowing the network to solve multiple tasks.

**Example on low rank dynamics:**
We are using dynamical mean field theory corresponding to large N limit, and we are using transfer function $\phi(x) = \tanh x$. The gain is $\phi'(x) = 1 - \tanh^2 x$.
For the rank 2 dynamics, we have the following dynamics

$$\frac{\tau d\boldsymbol{\kappa}}{dt} = -\boldsymbol{\kappa} + \boldsymbol{\kappa}\boldsymbol{\sigma_{mn}}\langle\phi'(0, \boldsymbol{\sigma_{m^2}}\boldsymbol{\kappa})\rangle$$

Just as an example, for special coupling pattern (ring attractor, or limit cycle):

$$\boldsymbol{\sigma_{mn}} = \begin{pmatrix} \sigma & -\sigma_w \\ \sigma_w & \sigma \end{pmatrix}$$

We have simplified equations in polar coordinate:

$$\tau\frac{d\rho}{dt} = -\rho + \rho\sigma\langle\phi'(0, \rho^2)\rangle$$

$$\tau\frac{d\theta}{dt} = \sigma_w\langle\phi'(0, \rho^2)\rangle$$

The relevant trajectory is shown in main text Figure 7.

For the rank 3 dynamics where we have:

$$\frac{d\kappa_{PFC}}{dt} = -\kappa_{PFC} + \tilde{\sigma}_{n_{PFC}m_{PFC}}\kappa_{PFC} + \tilde{\sigma}_{n_{PFC}m_{MD_1}}\kappa_{MD_1} + \tilde{\sigma}_{n_{PFC}m_{MD_2}}\kappa_{MD_2}$$
$$+\tilde{\sigma}_{n_{PFC}I_A}\nu_{I_A} + \tilde{\sigma}_{n_{PFC}I_B}\nu_{I_B} + \tilde{\sigma}_{n_{PFC}I_C}\nu_{I_C} + \tilde{\sigma}_{n_{PFC}I_D}\nu_{I_D}$$

$$\frac{d\kappa_{MD_1}}{dt} = -\kappa_{MD_1} + \tilde{\sigma}_{n_{MD_1}m_{PFC}}\kappa_{PFC} + \tilde{\sigma}_{n_{MD_1}m_{MD_2}}\kappa_{MD_2}$$

$$\frac{d\kappa_{MD_2}}{dt} = -\kappa_{MD_2} + \tilde{\sigma}_{n_{MD_2}m_{PFC}}\kappa_{PFC} + \tilde{\sigma}_{n_{MD_2}m_{MD_1}}\kappa_{MD_2}$$

The relevant circuit is in main text Figure 8. Here, $\kappa_{PFC}$, $\kappa_{MD_1}$ and $\kappa_{MD_2}$ are the latent variables of the three regions. The $\tilde{\sigma}_{n_{...}m_{...}}$ are the effective coupling, which is, for $p_{tot}$ populations (notice the $p_{tot}$ is different in different brain region, and we have 3 brain regions, including PFC, $MD_1$ and $MD_2$. For 3 population model, region of $MD_1$ and $MD_2$ both only contain 1 population):

$$\tilde{\sigma}_{mn} = \frac{1}{p_{tot}}\sum_{p=1}^{p_{tot}}\sigma_{mn}^{p}\,\langle\phi'\rangle_p$$

Here, $\langle\phi'\rangle_p$ (specifically for p-th population in PFC in 1st equation of the 3 coupled nonlinear ODE) is the corresponding to gaussian distribution with $0$ mean and variance:

$$\Delta^{\mathrm{p}} = \sqrt{(\sigma_{nm}^{p}\kappa_{PFC})^2 + \sum_{l=1}^{2}(\sigma_{n_{MD_l}m_{MD_l}}^{p}\kappa_{PFC})^2 + \sum_{q=1}^{4}(\sigma_{I_qI_q}^{p}\nu_q)^2}$$

Each population received input $\nu_q$. We then use our intuition from experiment to give value to all different $\tilde{\sigma}_{n_{...}m_{...}}$. For example, there are 4 populations of cells in PFC receiving 4 different input cues. cue 1 and cue 2 are correspond to first context, while cue 3 and cue 4 are correspond to first context. So one need to set different value for different populations. For example, the terms in $\tilde{\sigma}_{n_{PFC}m_{MD_1}}\kappa_{MD_1}$ need to be positive for population 1 and 2 because relevant cues for 1st context is amplified by 1st MD population, while the other terms need to be negative for population 3 and 4 because the 2nd context is suppressed by 2nd MD population.

**Capacity can be explained by embedding manifold**:
It has been discussed a lot [3] about how to embed the patterns into a neural network. Here is a simple derivation following simple dynamical system analysis. Consider the effective coupling weight $\boldsymbol{W}_{eff}$, and dynamics is simple ordinary differential equation (ODE):

$$\frac{d\boldsymbol{I}}{dt} = \frac{1}{\tau}(-\boldsymbol{I} + \boldsymbol{W}\boldsymbol{\phi}(I) + I_{ext}) = F(\boldsymbol{I})$$

We then consider the Jacobian:

$$\boldsymbol{J} = \frac{\delta \boldsymbol{F}}{\delta \boldsymbol{I}} = \frac{1}{\tau}(-\mathbf{1}_N + \boldsymbol{W_{eff}} diag(\frac{d\boldsymbol{\phi}}{d\boldsymbol{I}}))$$

If Jacobian have specific matrix factorization with singular value decomposition, and we denote $\Phi = diag(\frac{d\boldsymbol{\phi}}{d\boldsymbol{I}})$:

$$\boldsymbol{U\Sigma V} = \frac{1}{\tau}(-1_N + \boldsymbol{W_{eff}}\boldsymbol{\Phi})$$

Here, $U$ and $V$ are collections of vectors (row and column respectively) related to the task variables. That says, if total number of task variable is some small number $L \ll N$, then we have a low rank matrix factorization of Jacobian. Also, as long as we know what task variable is involved to create target dynamics, we can always know what the effective weight is $W_{eff}$ we want. We therefore know that the capacity is limited by the rank of matrix $U$ and $V$, which is $L$, with a maximum value $N$. This is consistent with early theory that the degree of the weight and sparsity of the weight is relevant parameter in neural network when we consider capacity.
